# A novel key player in cognitive striatal processes: the Pthlh interneurons

**DOI:** 10.64898/2025.12.02.691591

**Authors:** M. Díez-Salguero, L. Harder, M. Herr, M. Llorca-Torralba, E. Berrocoso, J. Hjerling-Leffler, A. B. Munoz-Manchado

## Abstract

Interneurons in the dorsal striatum provide essential modulatory control over the projecting neurons, known as medium spiny neurons (MSNs), regulating the output of this brain structure, which is critical for motor control, decision-making, and learning. The diversity of interneurons has been recently determined based on single-cell transcriptomic analyses identifying novel populations as the *Pthlh*-expressing interneurons, one of the most abundant and distinctive inhibitory populations in the human and mouse striatum. Here, we generated a novel Pthlh^cre^ knock-in mouse line that enables comprehensive study of this critical population. Quantitative histological analyses show that this line effectively targets the Pthlh interneuron class, which appears as a stable population across age, sex and the anterior-posterior axis. To elucidate their functional role, we performed selective chemogenetic inhibition of Pthlh interneurons, which caused significant impairments in striatum-dependent cognitive functions and reduced exploratory behaviorur. Importantly, selective silencing did not affect basal locomotion, motor coordination, or anxiety-like behavior. Monosynaptic rabies tracing revealed that Pthlh interneurons receive dense local input from MSN subtypes and diverse long-range afferents from thalamic and different cortical regions. We further confirm that this population displays electrophysiological heterogeneity encompassing fast-spiking and fast-spiking-like phenotypes. Overall, our findings establish Pthlh interneurons as a major inhibitory class essential for cognitive processing—but not gross motor function—acting as key integrators of local MSN feedback and thalamocortical signals. The novel Pthlh^cre^ mouse line provides a valuable resource for future studies of inhibitory diversity in basal ganglia function and related neurological disorders.

## Introduction

The basal ganglia are a set of interconnected subcortical nuclei that play essential roles in motivation, motor control, decision-making, and procedural learning [1–5]. Neural circuits within this system form part of the extrapyramidal motor network, and their dysfunction is implicated in several neurological and psychiatric conditions, including Parkinson’s disease, Huntington’s disease, obsessive–compulsive disorder, schizophrenia, and major depressive disorder [6–15]. Within the basal ganglia, the striatum—comprising the caudate nucleus and putamen in human—acts as the primary input structure, integrating glutamatergic afferents from the cortex and thalamus with dopaminergic projections from the midbrain [16–19]. Functionally, the striatum is divided into dorsal (DS), which is organized into dorsomedial (DMS) and dorsolateral (DLS) compartments that support goal-directed and habitual behaviours [20–26] and ventral striatum (nucleus accumbens).

Approximately 90–95% of striatal neurons are MSNs, which constitute the direct and indirect output pathways of the basal ganglia [27]. The remaining 5–10% are interneurons, classically subdivided into cholinergic and three GABAergic classes (fast-spiking (FS) parvalbumin (Pvalb) interneurons, low-threshold spiking somatostatin/neuropeptide Y/nitric oxide synthase (SST/NPY/nNos) interneurons, and calretinin interneurons that exhibit remarkable molecular, morphological, and functional diversity [28–31]. Despite their small numbers, interneurons exert powerful modulatory control over MSNs activity, finely tuning local microcircuit dynamics and overall basal ganglia output [32–35]. Single-cell/nucleus transcriptomics (scRNA-seq/snRNA-seq) has transformed our understanding of interneuron diversity in the striatum [36–44], identifying seven molecularly distinct interneuron populations in the mouse [39]—Npy/Sst, Npy/Mia, Cck/Vip, Cck, Chat, Th, and Pthlh— and eight in the human including a distinct *PVALB-*high transcriptomic class and the TAC3 population, the human homolog of the mouse *Th* class (described in the adult human brain in Garma, L. D. *et al.* [45] and also reported across different mammalian species [46, 47].

In the mice, Pthlh interneuron population is one of the most abundant population characterized by a continuum of *Pvalb* expression and graded electrophysiological properties along the dorsoventral axis, suggesting spatially organized input integration [39, 48, 49]. In the human DS PTHLH presents PVALB expression at low levels and is characterized by other markers as *OPN3* and *IL1RAPL2*. Interestingly PVALB population in humans constitute a distinct transcriptomic class, characterized by *PVALB* high expression among other set of markers as *GRIK3*. PTHLH population have also been identified in non-human primates [47], indicating that this population is broadly conserved across mammals and establishing PTHLH interneurons as a quantitatively significant and phylogenetically conserved component of the mammalian striatum.

At the molecular level, *PTHLH* encodes parathyroid hormone-like hormone also known as parathyroid hormone-related protein (PTHrP), a member of the parathyroid hormone (PTH) family that primarily signals through the PTH type 1 receptor (PTHR1), widely expressed throughout the human brain [50]. *Pthlh* regulates diverse developmental and physiological processes, including endochondral bone formation, epithelial–mesenchymal signaling, and calcium ion transport [51, 52]. Originally PTHLH was identified as a tumor-derived factor responsible for humoral hypercalcemia of malignancy [53], but it was described that PTHLH inhibits calcium channel activity in nervous system, exerting neuroprotective effects against excitotoxicity [54] while PTH increases intracellular calcium levels, promote apoptosis through calcium overload and facilitates the conversion of vitamin D into its active form, 1,25-dihydroxyvitamin D, which itself has neuroprotective and anti-inflammatory properties [55–57].

Abnormal PTH/PPTHLH signaling has been associated with neuronal calcium imbalance, neuroinflammation, and reduced cerebral perfusion, all of which are key mechanisms in neurodegenerative and psychiatric disorders [58, 59]. Patients with hyperparathyroidism often present cognitive impairment and depressive symptoms that improve after parathyroidectomy [60, 61]. Conversely, PTH deficiency has been linked to cytokine dysregulation and altered PTH secretion [62, 63]. These findings point toward a critical role of PTH/PTHLH signaling in neuronal calcium homeostasis, synaptic plasticity, and neuroprotection, with potential implications in cognitive and affective disorders [64–66].

Taken together, emerging evidence positions *PTHLH*-expressing interneurons as a key inhibitory population linking molecular, developmental, and functional aspects of striatal organization. Their unique abundance, evolutionary conservation, and potential involvement in calcium signaling pathways suggest that PTHLH interneurons may represent a critical cellular substrate through which endocrine and neuromodulatory signals influence basal ganglia circuitry. Functionally, PTHLH interneurons are expected to modulate MSN excitability and synchronization, contributing critically to the spatiotemporal organization of striatal microcircuits. Understanding their role in health and disease could provide new insights into the cellular mechanisms underlying motor, cognitive, and neuropsychiatric dysfunction.

### Material and Methods

#### Histological characterization

##### Transgenic Mice

For all experiments in this study, we used the knock-in Pthlh-Cre (Pthlh^cre^) mouse line on a C57BL/6J background (Cyagen, Germany), crossed or not with the reporter line B6.Cg-Gt(ROSA)26Sortm14(CAG-tdTomato)Hze/J (Pthlh-cre^rdTom^; <u>JAX #007914</u>), which exhibits robust tdTomato fluorescence following Cre-mediated recombination. Both male and female mice were used in all experiments. Mice were maintained under standard laboratory conditions (22 °C; 12:12 h light/dark cycle, lights on at 08:00 a.m.) with food and water available *ad libitum*.

##### Tissue Processing

For *in situ* hybridization, brains from Pthlh-cre^tdTomato^ mice were dissected, directly embedded in OCT compound (Tissue-Tek® O.C.T. Compound, Sakura), frozen on dry ice, and stored at −80 °C.

Brains for fluorescent immunohistochemistry (IHF) and combined fluorescent *in situ* hybridization–immunofluorescence (FISH–IHF) were obtained following transcardiac perfusion with phosphate-buffered saline (PBS) and 4% paraformaldehyde (PFA) using a perfusion pump at a flow rate of 12 ml/min. Brains were post-fixed in 4% PFA for 24 h, cryoprotected in 30% sucrose in PBS, and stored as described above.

Coronal brain sections (10 µm for histological characterization and DREADDs experiments, or 40 µm for rabies tracing) were obtained using a cryostat (Thermo Fisher Microm HM525) and collected on Superfrost Plus microscope slides. Slides were air-dried at room temperature (RT) for a few minutes, stored at −20 °C for 1 h, and transferred to −80 °C for long-term storage.

##### RNAscope high-sensitive fluorescent *in situ* hybridization (FISH)

High-sensitive *in situ* hybridization was performed using the RNAscope Multiplex Fluorescent Reagent Kit v2 (323110, Advanced Cell Diagnostics) on striatal sections from Pthlh-cre^tdTomato^ mice. Brains from three males at postnatal days (P) 10, 15, 21, 60–80, and 100–150, and three females at P21 and P100–150 were processed to detect single mRNA molecules.

Experiments were carried out following the manufacturer’s instructions (UM 323100, Advanced Cell Diagnostics) with minor modifications in tissue pretreatment where we treat the sections with Protease III for 5 min RT to improve antigen accessibility. The following probes were used: *Pthlh* (456521), *tdTomato* (317041-C2), *Pvalb* (421931-C3), and *Cdh5* (312531-C3). Detection was performed using TSA Vivid 520 (323271), TSA Vivid 570 (323272), and TSA Vivid 650 (323273) at 1:1500 (Advanced Cell Diagnostics).

Before mounting in Fluoromount-G (0100-01, SouthernBiotech), nuclei were counterstained with DAPI. All experiments were completed within a one-day protocol to ensure optimal tissue preservation and signal quality.

##### RNAscope high-sensitive fluorescent *in situ* hybridization-immunofluorescence (FISH-IHF)

High-sensitive FISH was performed using the RNAscope Multiplex Fluorescent Reagent Kit v2 as previously described, with minor modifications in the pretreatment steps. Slides were first air-dried, and a hydrophobic barrier was drawn around selected sections. Sections were then incubated in target retrieval solution for 5 min at 99°C on a hot plate, rinsed twice in distilled water (2 min each), and dehydrated twice in 100% ethanol (2 min each RT). After air drying for 5 min, sections were treated with Protease IV for 20 min RT, followed by two washes in distilled water (2 min each).

RNA hybridization were performed using the following probes: *Pthlh* (456521), *Pvalb* (421931-C3), *Th* (317621-C3), *Cck* (402271-C2), *Chat* (408731-C3), *Sst* (404631-C3), *Ppp1r1b* (405901-C2), *Drd1a* (406491), and *Drd2* (406501). After signal detection with TSA Vivid reagents for each channel, immunohistochemistry was performed.

Sections were washed in PBS and incubated in blocking solution (10% normal goat serum, 2.5% BSA, 0.5 M NaCl, and 0.3% Tween 20 in PBS) for 30 min RT. They were then incubated overnight RTwith primary antibodies diluted in dilution buffer (2.5% BSA, 0.5 M NaCl, and 0.3% Tween 20 in PBS), followed by three washes in 0.05% Tween 20 in PBS. Secondary antibodies were applied for 2 h RT and then washed three times in 0.3% Tween 20 in PBS. Nuclear counterstaining was performed using DAPI (1:500 in water, 2 min), followed by one final wash in 0.3% Tween 20 in PBS and mounting with Fluoromount-G.

Primary antibodies were used at the following concentrations: chicken anti-GFP (1:200; Abcam, ab13970) and rabbit anti-DsRed (1:100; Takara, 632496). Secondary antibodies were goat anti-chicken Alexa Fluor 488 (1:500; Invitrogen, A-11039) and goat anti-rabbit Alexa Fluor 555 (1:500; Invitrogen, A-21428).

##### Fluorescent immunohistochemistry

Sections were washed twice in PBS for 15 min each, fixed in 4% PFA for 5 min, and then washed three times in 0.1% Tween-20 in PBS. Blocking was performed for 1 h RT in a solution containing 5% normal goat serum, 2.5% BSA, 0.5 M NaCl, and 0.3% Tween-20 in PBS.

Sections were then incubated overnight at 4 °C with primary antibodies diluted in dilution. After four washes in 0.1% Tween-20 in PBS (15 min each), sections were incubated with secondary antibodies for 4 h RT, followed by three additional washes in 0.1% Tween-20 in PBS (10 min each).

Nuclear counterstaining was performed with DAPI (1:5000 in water) for 5 min. Sections were then rinsed once in PBS before mounting.

Primary antibodies concentrations:

- Chicken anti-GFP (1:2000; Abcam, ab13970)
- Rabbit anti-DsRed (1:300; Takara, 632496)

Secondary antibodies concentrations:

- Goat anti-chicken Alexa Fluor 488 (1:1000; Invitrogen, A-11039)
- Goat anti-rabbit Alexa Fluor 555 (1:1000; Invitrogen, A-21428) <u>Image acquisition:</u>

Confocal imaging was performed using a Zeiss LSM900 microscope equipped with an Airyscan detector and controlled by ZEN 2.6 software (Zeiss, Germany). To capture the entire DS or whole coronal sections, tile scans were acquired using a 20× air/dry objective. Final images were automatically stitched using the integrated stitching function in the ZEN software.

##### Image analysis

Quantitative image analysis was performed using QuPath software (version 0.4.3) following this workflow: (1) Create training images for each channel using enough selected sections per staining. Within each section, a square region of interest (ROI) of approximately 600,000 µm² was drawn. (2) Cell and subcellular detection parameters were optimized for each staining and applied to the training images for RNA molecules detection. (3) An object classifier for each marker was trained by manually labeling positive cells. Minimum cut-off values for the number of subcellular spots were defined per marker and applied manually (tdTomato > 15the cut-off wa and for remaining genes ≥ 5 estimated spots) to consider a cell positive. (4) Individual classifiers were combined into a composite classifier. (5) ROIs were defined across the DS based on DAPI staining but excluding artifacts and high fluorescent vessels. (6) The adjusted detection settings and composite classifier were applied to all ROIs for each staining. (5) The resulting list of cells and assigned markers per ROI was exported for evaluation. Unexpected marker combinations or abnormal spot counts were manually verified and corrected if necessary.

#### Electrophysiology

##### Slice electrophysiology

Whole-cell patch clamp recordings were acquired of tdTomato-positive cells in acute brain sections prepared from Pthlh^tdTomato^ mice between P70-130 of both sexes. Animals where anaesthetized deeply with isoflurane and ketamine/xylazine (4:1), followed by transcardial perfusion with ice cold oxygenated cutting solution (in mM: 62.5 NaCl, 100 sucrose, 2.5 KCl, 25 NaHCO3, 1.25 NaH2PO4, 7 MgSO4, 0.5 CaCl2, and 10 glucose). The brain was immediately dissected and cut into 300µm sections at a speed of 0.12mm/s using a vibratome (Leica, VT 1200S) while being submerged in ice cold oxygenated cutting solution continuously. Single slices were immediately transferred to a section holder with oxygenated artificial cerebrospinal fluid (aCSF) solution (in mM: 124 NaCl, 2.5 KCl, 26 NaHCO3, 1.2 NaH2PO4, 2 MgSO4, 2 CaCl2, and 20 glucose). Slices were incubated at 35°C for 30 minutes and afterwards kept RT. During recording, slices were continuously perfused with oxygenated aCSF. Patch pipettes were pulled from borosilicate glass capillaries (Hilgenberg, #1403512) to a resistance of 4-8 MΩ and filled with intracellular solution (in mM: 150 K-gluconate, 3 MgCl2, 0.5 EGTA, 10 HEPES, 2 MgATP, 0.3 Na2GTP). The signal was amplified with Multiclamp 700B (Molecular Devices) and digitized with Digidata 1550 (Molecular Devices).

### Measurement and analysis of intrinsic electrophysiological properties

In current-clamp mode, 1pA depolarizing current steps were applied to isolate the first action potential (AP) discharge. The following electrical properties were extracted from this threshold AP, using the python packages ipfx and feel [67, 68]: AP Half width, AP threshold, AP latency, rheobase, AP amplitude, AHP amplitude, and AHP latency.

Input resistance, inter-spike-interval, and the time constant were measured using Clampfit (version 10.7, Molecular Devices). Input resistance was measured by using the quick graph function in Clampfit to generate an I-V graph on subthreshold recordings. The slope of a linear regression line represented the input resistance. Inter-spike-interval was measured on the first 100ms of the highest frequency sweep before failure. The time constant was measured from a hyperpolarizing current step (causing a hyperpolarization between -90 to -100mV) by applying an exponential fit to the decay phase of the voltage response.

The antero-posterior location of each cell (in µm from bregma) was determined by inspection of the respective slice (10x objective) and determining the most similar section in the Allen brain atlas (Allen Brain Reference Atlas, Mouse P56 Coronal). The medio-lateral and dorso-ventral localization of each cell (in µm) was estimated using the scaling factor of the respective Allen brain atlas image.

Clustering of cells was performed as described previously [48] using R. Principal component analysis was performed on half width and maximum frequency followed by k-means clustering.

#### Surgeries

Mice (8-week-old) were anesthetized with 2% isoflurane using a low-flow vaporizer system (SomnoFlo, Kent Scientific), mounted on a stereotaxic apparatus (World Precision Instruments), and maintained under 1–1.5% isoflurane throughout the surgery. For infusions, pulled glass capillary micropipettes (World Precision Instruments) were backfilled with the solution and injections were performed at a rate of 150 nL/min using a precision injection system (Nanoliter 2020, World Precision Instruments). The micropipette was left in place for at least 10 min following each injection before being slowly withdrawn. Injections were performed bilaterally at the following coordinates (relative to bregma): +0.61 mm AP, ±1.75 mm ML, and −3.5 mm DV. Carprofen (5 mg/kg) was administered subcutaneously for post-operative analgesia over 24–48 h, including the day of surgery.

For Cre-dependent hM4D(Gi)-GFP expression in the DS, 500nL of pAAV8-hSyn-DIO-hM4D(Gi)-EGFP (3.5 × 10¹² vg/mL; Viral Vector Core, Duke University School of Medicine, Durham) or pAAV8-hSyn-DIO-EGFP (5.6 × 10¹² vg/mL; Viral Vector Core, Duke University School of Medicine, Durham) for control animals was injected into the striatum of Pthlh-cre^tdTomato^ mice.. To inactivate hM4D(Gi)-expressing neurons, Clozapine N-oxide (CNO, 4 mg/kg; Biosynth FC20527) dissolved in saline was administered intraperitoneally 30 min before each behavioural session. Control mice received an equivalent volume of saline.

For rabies tracing, 100 nL of pAAV8-hSyn-TVA66T-G-mCherry (4.75 × 10¹² vg/mL; Kavli Institute for Systems Neuroscience, Norway) or pAAV8-hSyn-TVA-G-mCherry (5.97 × 10¹² vg/mL; Kavli Institute for Systems Neuroscience, Norway) was injected into the DS Pthlh^cre^ mice. Three weeks later, 500 nL of EnvA-pseudotyped GFP-expressing RABV (5.0 × 10⁸ cfu/mL; Kavli Institute for Systems Neuroscience, Norway) was injected into the same site. Mice were sacrificed five days later.

For retrograde tracing, red fluorescent microspheres (Lumafluor) were diluted 1:10 in saline, and 500 nL of the tracer was infused into the dorsal striatum of Pthlh^cre^ mice using the same procedure as described above. After a suitable survival period for retrograde labeling, animals were sacrificed for histological analysis.

#### Data Analysis for Whole-Brain Tracing

The regions selected for analysis included the major local and long-range inputs to the striatum. For local inputs, striatal sections near and within the injection site from brains infused with the TVA^66T^ helper virus were analyzed. GFP-positive cells, co-labeled or not with additional markers, were quantified using QuPath software. The number of labeled neurons in each section was normalized to the total number of input neurons detected within the striatum.

For long-range inputs, GFP-positive neurons from brains injected with the conventional TVA helper virus were counted across serial sections throughout the entire brain, excluding striatal sections. Because the total number of input neurons varied between brains, cell counts for each region were normalized to the total number of input neurons detected in the same brain [69, 70].

The average number of input neurons across striatal regions in TVA^66T^ brains and the average number of input neurons across distant regions in TVA brains were used to calculate the relative percentage of local versus long-range inputs.

#### Behavioural Experiments

Behavioural experiments were conducted in dedicated testing rooms during the light phase (09:00–15:00 h). Mice were acclimated in an adjacent holding room for at least 1h prior to testing. All apparatuses were thoroughly cleaned with 70% ethanol between trials to eliminate olfactory cues. Animals underwent multiple behavioural assays with sufficient rest between tests, following an ascending order of stress intensity to minimize carryover effects.

#### Novel Object Recognition (NOR) Test

Novel object recognition short-term (STM) and long-term memory (LTM) was assessed. Mice were tested in a 45 × 45 cm square arena with opaque walls, using plastic objects differing in shape, color, and texture. Animals were habituated to the empty arena for 15 min the day prior to testing. On the test day, mice underwent a 5 min habituation phase in the absence of objects, followed by a 15 min training phase in which two identical objects (A and A’) were placed in opposite corners. 90 min (STM, as learning index) and 24 h (LTM, as memory index), after training session, mice were reintroduced to the arena for a 10 min test in which one familiar object was replaced by a novel one (B for STM, C for LTM). Object exploration was defined as actively sniffing or touching the object while maintaining gaze; circling or sitting on the object was excluded. Exploration was quantified as the number of interactions, percentage preference, and a Discrimination Index (DI), calculated as DI = (T_novel – T_familiar) / (T_novel + T_familiar) [71].

#### Elevated Plus Maze (EPM) Test

Each mouse was placed in a gray, cross-shaped maze (Panlab, S.L., Barcelona, Spain) elevated 40 cm above the floor, consisting of two open arms (29.5 × 6 cm), two closed arms (29.5 × 6 × 15 cm with walls), and a central square (6 × 6 cm). Lighting in the central square was maintained at 11 lux. Behaviour was recorded for 5 min, and the percentage of time spent in the open arms was quantified using SMART video-tracking software as an index of anxiety-like behaviour [72].

#### Rotarod Test

Motor coordination and balance were evaluated using a rotarod apparatus (Panlab, S.L., Barcelona, Spain). Mice were tested under two conditions: constant speed (5 rpm) for up to 5 min, conducted in three trials with 15 min rest between trials, and accelerated speed (4–40 rpm over 5 min), conducted in a single trial [73]. The number of falls was recorded manually, and latency to fall was automatically measured by the apparatus.

#### Hole Board Test

Mice were placed in the center of a plastic arena (50 × 50 × 35 cm; Panlab, S.L., Barcelona, Spain) containing 16 holes (4 cm diameter). On the first day, animals explored the hole board for 6 min during a training phase in which all holes were empty. On the second day, identical objects were placed in 8 of the holes, with the remaining 8 left empty. Mice were allowed to explore for 6 min, during which locomotor activity, as well as the location, number, and duration of hole visits, were recorded [74].

#### Open Field Test (OFT)

Each mouse was placed in the center of a circular arena (45 cm diameter) enclosed by continuous opaque walls, under constant dim lighting (10–15 lux), and allowed to explore freely for 20 min while being recorded. The central area was defined as a circle covering approximately 50% of the total arena. Time spent in the center, total distance traveled, and the frequency and duration of rearings were quantified using SMART video-tracking software (Panlab, S.L., Barcelona, Spain) [75].

#### Statistical Analysis

Data are presented as mean ± SEM. Statistical analyses were performed using GraphPad Prism 9.5.1 (GraphPad, San Diego, CA). Grubbs’ test was used to identify statistical outliers and the Shapiro-Wilk test to evaluate the normality of data distribution. Comparison between two groups were conducted using unpaired Student’s t tests or Wilkoxon test for paired without normal distribution measurements. For multi-group or multi-factor analyses, one- or two-way ANOVA, with or without repeated measures, was applied, followed by Tukey’s, Kruskal-Wallis or Dunnett’s post hoc tests, as appropriate. Correlations were assessed using Pearson’s correlation. Statistical significance was set at p < 0.05.

## Results

We have generated a Pthlh^cre^ knock-in mouse to investigate the recently described Pthlh interneuron population^51,56,58,59,85,86^. Using this mouse line we have: (1) histologically characterized the Pthlh interneuron population using FISH; (2) electrophysiologically identified distinct subtypes within the Pthlh population; (3) studied the functional role of Pthlh interneurons through chemogenetic inhibition of *Pthlh*-expressing cells in the DS; and (4) analyzed both local and global connectivity patterns of *Pthlh*-expressing cells in the DS (Figure 1A).

**Figure 1.**
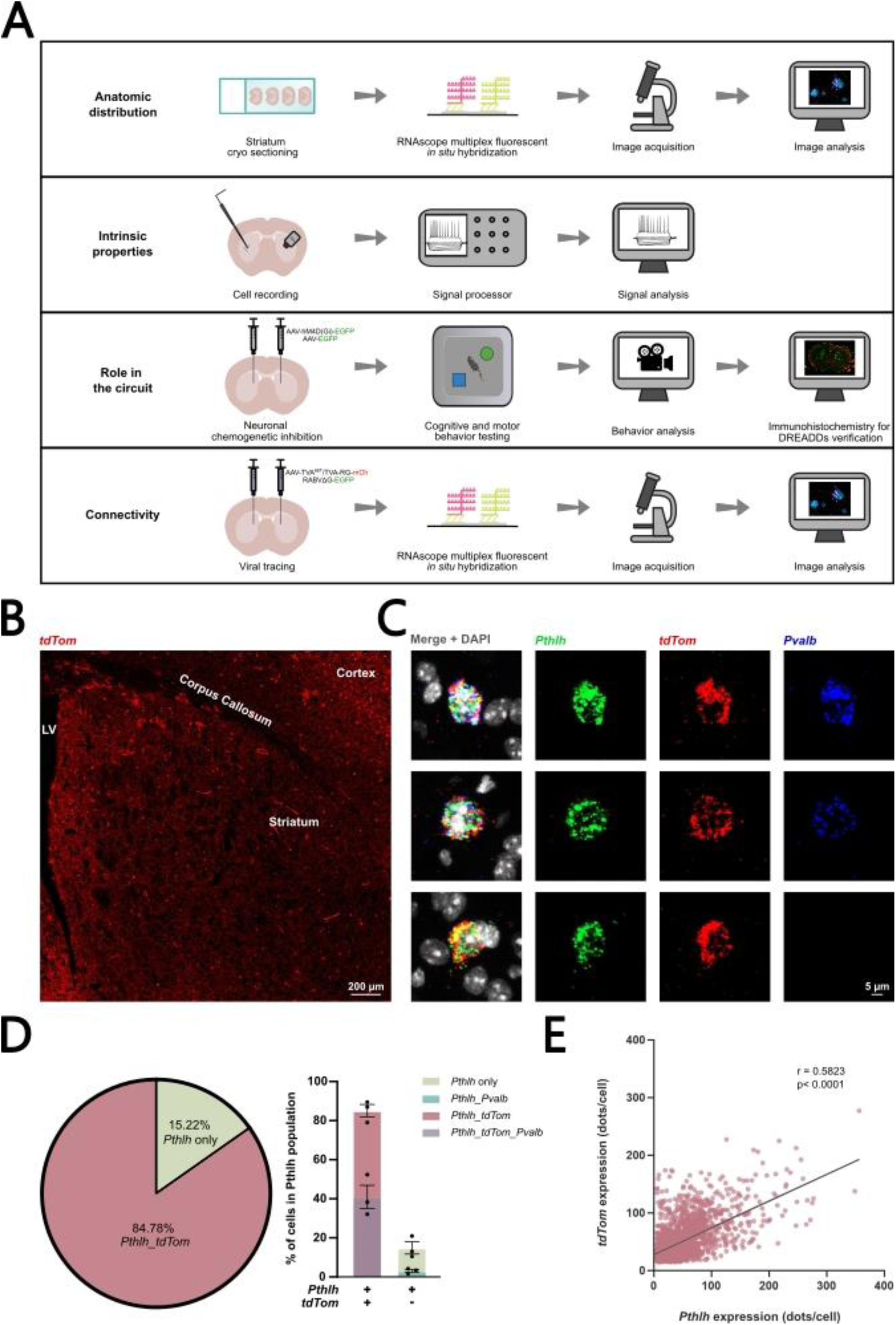
Validation by quantitative fluorescence *in situ* hybridization confirms the labelling of *Pthlh*-expressing cells in adult male mice of the Pthlhtd^Tomato^ line. A. Schematic overview of experimental design. **B.** Immunohistochemistry for tdTomato (tdTom) showing the reporter-labeled cells in a dorsal striatum of the Pthlh^tdTomato^ mouse line. **C.** Fluorescent *in situ* hybridization for *Pthlh*, *tdTomato* and *Pvalb* illustrating double- and triple-positive cells within the Pthlh population, displaying high (top), medium (middle), or no (bottom) *Pvalb* expression. **D.** Pieplot (left) and barplot (right) about the Pthlh population coexpressing *tdTomato* and *Pvalb* (n = 3). **E.** Scatter plot showing the correlation of expression levels (dots/cell) between *Pthlh* and *tdTomato* (n = 3; 1 776 cells). Error bars represent mean ± SEM. r and p values were acquired using Pearson’s correlation.

### 1. The knock-in Pthlh^cre^ mouse line allows to target most Pthlh interneurons

To validate the Pthlh^cre^ mouse line, we crossed it with the reporter line B6.Cg-Gt (ROSA)26Sor^tm14(CAG-tdTomato)Hze^/J, generating the Pthlh-cre^tdTomato^ mouse line (Figure 1B). Mice of different ages (P10, P15, P21, P60-80 and P100-150 days) and both sexes were used to perform high-sensitive FISH using probes against *Pthlh*, *tdTomato*, *Pvab* and *Cdh5* (Figure 1C, Supplementary Figure 1E). We quantified the number of cells expressing each gene and their possible combinations. The analysis revealed that 85.14 ± 5.45% of the *Pthlh-*expressing cells were labeled by the reporter gene *tdTomato* and only 14.86 ± 5.45% of *Pthlh+ c*ells were not labeled by the reporter in adult male mice (P100-150 days). Among them 44.23 ± 9.36% were Pthlh+/tdTomato+/Pvalb-, 40.91 ± 10.38% Pthlh+/tdTomato+/Pvalb+, 11.78 ± 4.33% Pthlh^+^/tdTomato^-^/Pvalb^-^ and the 3.08 ± 1.33% were Pthlh^+^/tdTomato^-^/Pvalb^+^ (Figure 1D). There is a positive correlation observed between the expression level of *Pthlh* and *tdTomato* (Figure 1E). Focusing on the reporter-labeled population the 64.96 ± 7.24% were *Pthlh-*expressing cells. By including *Pvalb* expression we found that 34.29 ± 10.90% cells were tdTomato^+^/Pthlh^+^/Pvalb^-^ and 30.67 ± 4.03% were tdTomato^+^/Pthlh^+^/Pvalb^+^). The 35.04 ± 7.24% of *tdTomato*⁺ cells did not express *Pthlh* in adult mice, were the 33.91 ± 6.95% of cells are tdTomato^+^/Pthlh^-^/Pvalb^-^ and the remaining 1.13 ± 0.30% were tdTomato^+^/Pthlh^-^/Pvalb^+^ (Supplementary Figure 1C). Among these non-*Pthlh tdTomato*⁺ cells, 19.75 ± 2.71% were endothelial cells. Notably, only 6.68 ± 0.82% of *Cdh5*-positive endothelial cells co-expressed the reporter in adult male mice (Supplementary Figure 1E, F–G). No significant sex- or age-related differences were observed in the percentage of *Pthlh* cells labeled by the reporter, or in the proportion of reporter-labeled cells that were *Pthlh*-expressing (Supplementary Figure 1A, B).

### 2. Consistent abundance of Pthlh interneurons along the rostro-caudal striatum with age-related increase in *Pvalb* expression

We aimed to identify the Pthlh distribution along the rostro-caudal and dorso-ventral axis. To do that we have characterized the Pthlh interneuron population by performing FISH against *Pthlh* and *Pvalb* across different groups of age and sex Pthlh-cre^tdTomato^ mice described above, and at multiple levels along the rostro-caudal axis of the DS (Figure 2A).

**Figure 2.**
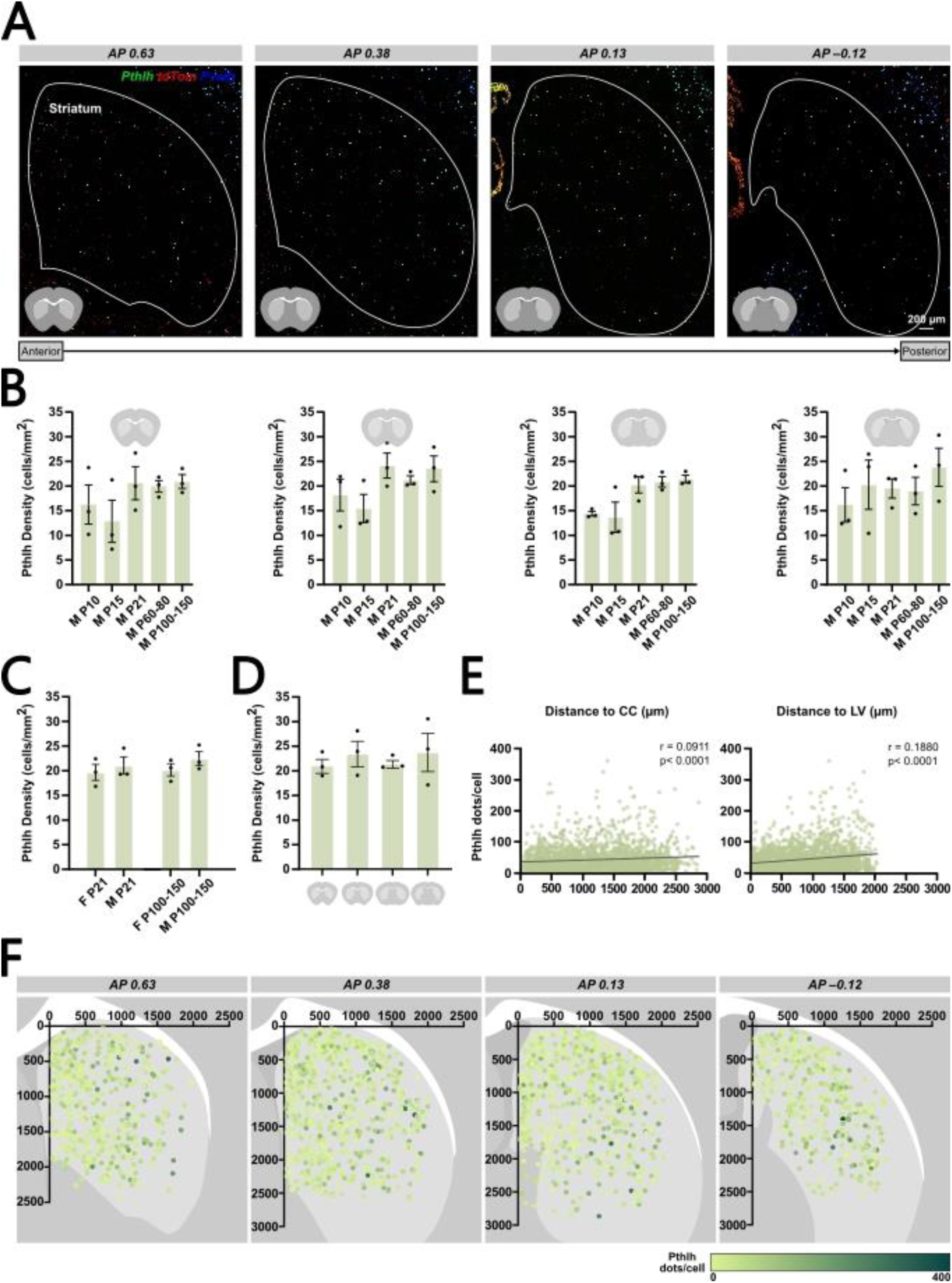
Histological characterization of the *Pthlh*-expressing interneurons in the dorsal striatum. **A.** Fluorescent *in situ* hybridization for *Pthlh*, *tdTomato* and *Pvalb*, along the rostro-caudal axis at four stereotaxic levels: Bregma +0.63 ± 0.05, +0.38 ± 0.05, +0.13 ± 0.05, and –0.12 ± 0.05 mm. **B.** Barplot showing the density of *Pthlh*-expressing cells across different experimental groups and rostro-caudal levels (n = 3 per group). **C.** Same as in B but comparing females and males at P21 and P100-150 (n = 3 per group). **D.** Bar plot showing the rostro-caudal distribution of *Pthlh*-expressing cells in P100–150 male mice (n = 3). **E.** Scatter plots showing the correlation between *Pthlh* expression levels (dots per cell) and anatomical position along the dorsoventral (right) and mediolateral (left) axes in P100–150 male mice (n = 3; 2 008 cells). **F.** Schematic representation of the spatial distribution and expression levels of *Pthlh*-expressing cells in the dorsal striatum of P100–150 male mice, across the same rostro-caudal levels as in panel A (n = 3; 2 008 cells). Error bars represent mean ± SEM. r and p values were acquired using Pearson’s correlation.

Quantification of the *Pthlh*-expressing population revealed a density of 22.45 ± 2.48 cells/mm² in adult male mice, with no significant differences observed along the rostro-caudal axis (Figure 2D). Similarly, no significant differences in Pthlh cell density in DS were found across age groups—P10 (16.32 ± 4.62), P15 (15.54 ± 4.99), P21 (21.05 ± 3.04), and P60–80 (20.17 ± 0.88)—or between sexes at P21 (females: 19.64 ± 2.80; males: 21.05 ± 3.04) and at P100–150 (females: 20.18 ± 2.17; males: 22.45 ± 2.48) (Figure 2C, Supplementary Figure 2A). No rostro-caudal differences were detected in any group (Figure 2B, Supplementary Figure 2A). While the number of *Pthlh*-expressing cells remained stable across age and along the rostro-caudal axis, the number of *Pvalb* expressing cells within the Pthlh population in the DS increased with age. Their density in P10, P15, P21, P60–80, and P100–150 days mice was 5.07 ± 1.26, 8.41 ± 4.60, 12.24 ± 1.73, 11.67 ± 2.06, and 10.78 ± 1.94 cells/mm², respectively (Supplementary Figure 2A, D). No significant sex differences were observed at P21 (females: 10.59 ± 2.12; males: 12.24 ± 1.73) or at P100–150 (females: 12.25 ± 1.13; males: 10.78 ± 1.94) days in the DS (Supplementary Figure 2E). Furthermore, Pvalb cell density within the Pthlh population remained stable along the rostro-caudal axis in adult male mice (Supplementary Figure 2B, F).

Quantitative high-sensitive FISH revealed a very weak correlation between *Pthlh* expression and the distance to the corpus callosum (dorsoventral axis: r = 0.0911, p < 0.0001) and to the lateral ventricle (mediolateral axis: r = 0.1880, p < 0.0001) (Figure 2E, F). In contrast, *Pvalb* expression showed a stronger spatial correlation, with higher expression observed in Pthlh cells located in the ventrolateral region of the DS (dorsoventral axis: r = 0.1873, p < 0.0001; mediolateral axis: r = 0.3888, p < 0.0001) (Supplementary Figure 2G, H). Additionally, a positive correlation was found between *Pthlh* and *Pvalb* expression levels within the *Pthlh*-expressing population (r = 0.6107, p < 0.0001) (Supplementary Figure 2C).

### 3. Electrophysiological diversity of Pthlh interneurons revealed by the Pthlh^cre^ mouse line

To investigate the electrophysiological properties and diversity within the Pthlh population, we electrophysiologically recorded 59 cells from 20 Pthlh^tdTomato^ mice. These cells were classified based on their electrophysiological profiles as FS (12 cells, 20.34%), FSL (28 cells, 47.46%), MSN (5 cells, 8.47%), and non-FS (5 cells, 8.47%). Nine cells were excluded from analysis (Figure 3A). To differentiate Pthlh-recorded cells into FS or FSL, we performed principal component analysis and k-means clustering on AP half-width and maximum frequency —two parameters that have been shown to best distinguish these populations (Figure 3B). We observed that FS and FSL cells were distributed throughout the DS with no apparent spatial bias (Figure 3C). Several electrophysiological parameters differed significantly between the FS and FSL groups: AP amplitude (mV) and maximum frequency (Hz) were significantly lower, while AP threshold (mV), half-width (ms), and AHP latency (ms) were significantly higher in FSL cells compared to FS cells (Figure 3D).

**Figure 3.**
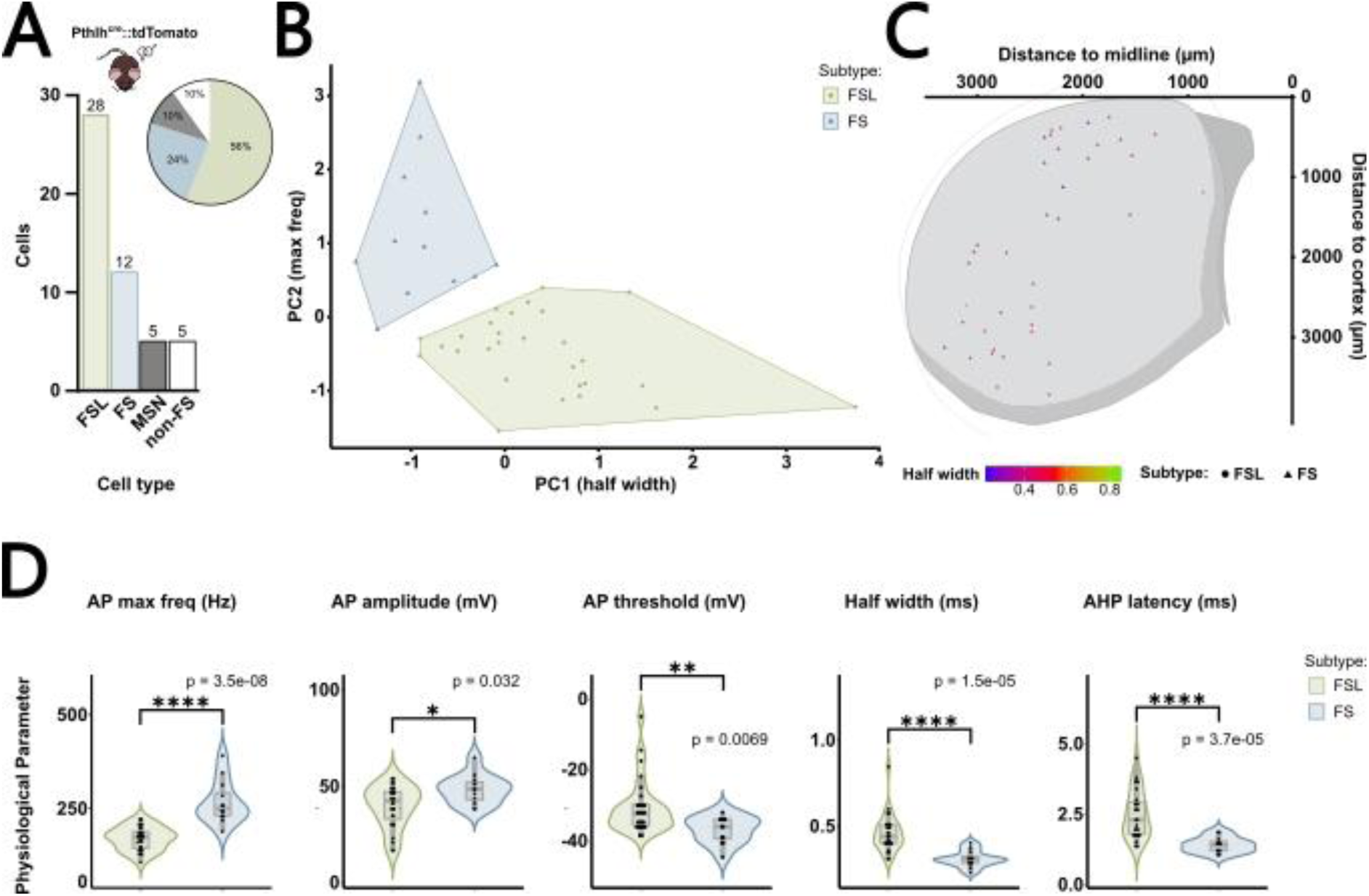
Electrophysiological properties within the Pthlh interneuron population. **A** Barplot showing the number and proportion of different electrophysiological profiles recorded (n = 20 mice, 50 cells). **B.** Scatter plot displaying clusters based on maximum firing frequency (Y-axis) and half-width (X-axis). **C.** Scatter plot showing the spatial distribution of fast-spiking (FS, triangles) and fast-spiking-like (FSL, circles) cells along the dorsoventral (Y-axis) and mediolateral (X-axis) axes colored by half-width. **D.** Comparison of electrophysiological parameters between striatal FS (n = 12) and FSL (n = 28) cells. Statistical significance was assessed using Wilcoxon rank sum test. Whiskers represent 1.5 * IQR.

Together, these results are consistent with the previously reported electrophysiological heterogeneity within the Pthlh population^51,57^, and indicate that our genetic strategy captures nearly the entire Pthlh population, unlike previous studies where this was not achievable.

### 4. Pthlh interneurons are involved in cognitive processes

We performed selective chemogenetic inhibition of *Pthlh*-expressing cells using hM4D(Gi)-DREADDs [76–78] (Figure 4A–E) followed by comprehensive battery of behavioural tests covering motor (rotarod and open field, OF) and cognitive (new object recognition, NOR, and hole board test, HB) aspects as well as anxiety (elevated plus maze, EPM) to investigate the role of *Pthlh*-expressing interneurons in the circuit (Figure 4A).

**Figure 4.**
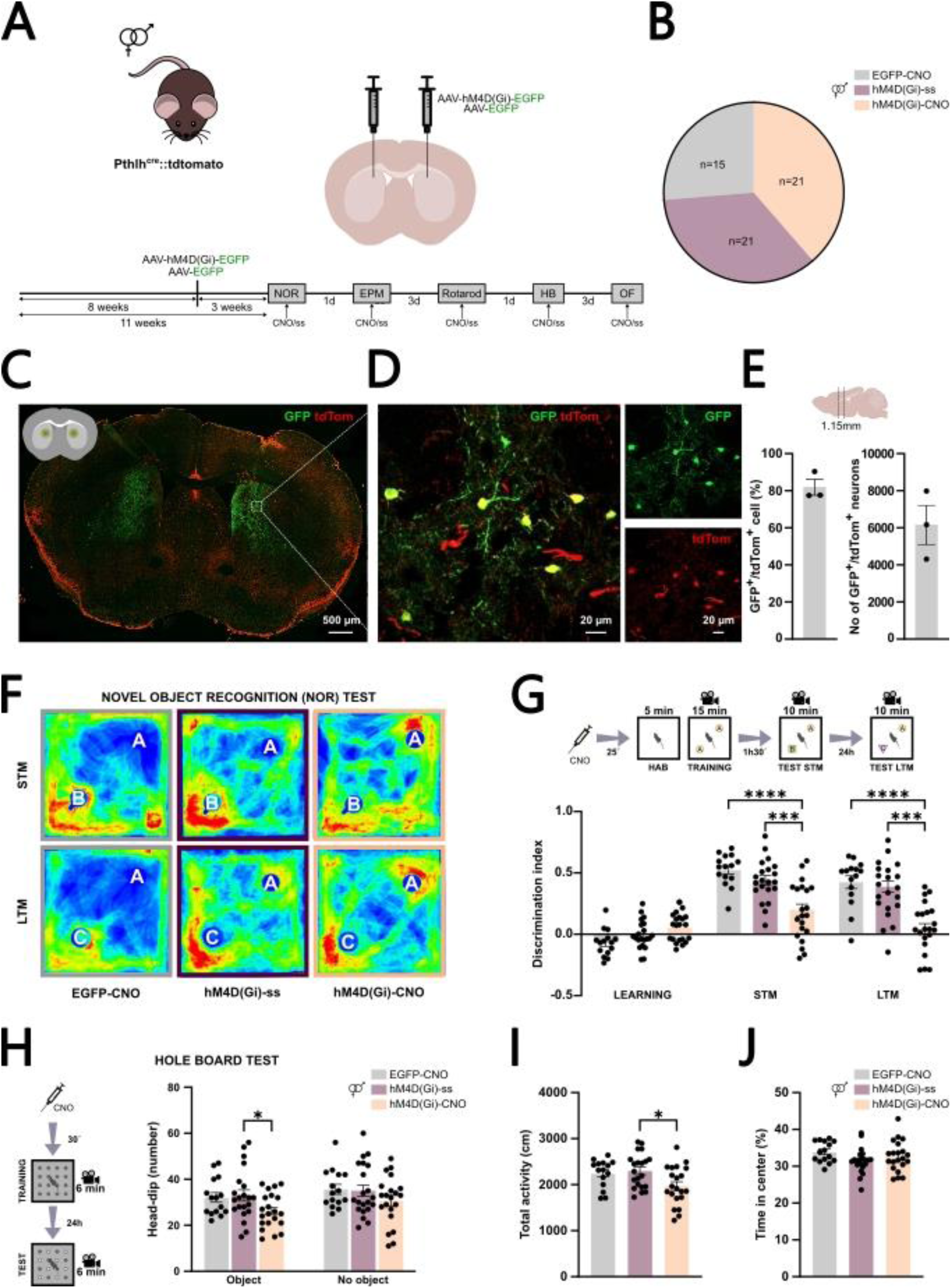
Cognitive and exploratory impairments caused by selective inhibition of *Pthlh*-expressing cells in the dorsal striatum. **A** Schematic representation of experimental design. **B.** Pieplot showing the number of animals in each experimental group: EGFP-CNO (mice injected with the control virus and administered CNO 30 minutes before the behavioral session), hM4D(Gi)-ss (mice expressing the inhibitory DREADD and administered saline 30 minutes before the behavioral session), and hM4D(Gi)-CNO (mice expressing the inhibitory DREADD and administered CNO 30 minutes before the behavioral session). **C.** Representative immunohistochemistry image showing *Pthlh*-expressing cells (tdTomato) infected with the inhibitory hM4D(Gi)-DREADD virus (GFP). **D.** Magnified view from panel C. **E.** Percentage of infected cells that are Pthlh (left; n = 3) and total number of Pthlh infected cells in the striatum (right; N = 3). **F.** Representative heatmaps showing exploration around the objects. Familiar objects are labeled as A; novel objects for short-term memory (STM) and long-term memory (LTM) are labeled as B and C, respectively. **G.** Schematic of the Novel Object Recognition (NOR) protocol (top) and discrimination index after learning, STM, and LTM phases (bottom). **H**. Schematic representation of Hole board test protocol (left) and barplot showing the number of head-dips in holes with and without objects per group (right). **I.** Barplot showing total activity levels across groups. **J.** Barplot showing the percentage of time spent in the center of the arena per group. Error bars represent mean ± SEM. * p<0.05, *** p<0.001 and **** p<0.0001, assessed by one-way ANOVA with repeated measures followed by Tukey’s or Kruskal-Wallis post hoc test.

Interestingly, no differences were found in any of the motor applied tests. These motor abilities were evaluated following Pthlh interneuron silencing. In the rotarod test mice were subjected to three trials at a constant speed of 5 rpm, followed by one accelerating trial where the speed increased from 4 to 40 rpm over 5 minutes. No significant differences were observed between experimental and control groups in any of the constant-speed trials. All groups showed improved performance over sessions, with fewer falls in the last trial (hM4D(Gi)-CNO: 0.71 ± 0.22; hM4D(Gi)-ss: 0.71 ± 0.14; EGFP-CNO: 1.00 ± 0.34) compared to the first (hM4D(Gi)-CNO: 2.38 ± 0.43; hM4D(Gi)-ss: 2.14 ± 0.45; EGFP-CNO: 3.73 ± 0.49) (Supplementary Figure 3J). Similarly, in the accelerating rotarod test (4–40 rpm), no group differences were detected in the latency to fall (hM4D(Gi)-CNO: 57.74 ± 6.15; hM4D(Gi)-ss: 68.80 ± 6.86; EGFP-CNO: 46.60 ± 7.47) (Supplementary Figure 3K).

Consistently, the OF test showed no differences in total distance travelled between groups (hM4D(Gi)-CNO: 5 420 ± 211.3; hM4D(Gi)-ss: 5 775 ± 341.2; EGFP-CNO: 6 049 ± 263.1) (Supplementary Figure 3H, I). Likewise, the percentage of time spent in the center of the arena was comparable across conditions (hM4D(Gi)-CNO: 22.74 ± 1.61; hM4D(Gi)-ss: 23.89 ± 1.74; EGFP-CNO: 20.93 ± 2.26) (Supplementary Figure 3D, E). The number and duration of rearing events also did not differ between groups (number: hM4D(Gi)-CNO: 77.71 ± 9.78; hM4D(Gi)-ss: 86.37 ± 11.10; EGFP-CNO: 79.93 ± 8.90; duration: hM4D(Gi)-CNO: 43.15 ± 7.24; hM4D(Gi)-ss: 46.04 ± 6.76; EGFP-CNO: 46.60 ± 6.79) (Supplementary Figure 3F, G).

Among the behavioural cognitive tests, we used the NOR to assess the potential impact of Pthlh interneuron inhibition on learning and memory. During the 15-minute training phase, all groups showed similar discrimination indices while exploring two identical objects (hM4D(Gi)-CNO: 0.053 ± 0.024; hM4D(Gi)-ss: -0.004 ± 0.025; EGFP-CNO: -0.069 ± 0.028) (Figure 4G). However, in the test phase for short-term memory (STM) mice with inhibited Pthlh interneurons failed to distinguish the new object and displayed significantly lower discrimination indices compared to controls (hM4D(Gi)-CNO: 0.187 ± 0.051; hM4D(Gi)-ss: 0.437 ± 0.037; EGFP-CNO: 0.527 ± 0.034) (Figure 4F, G). Similar results were observed in the long-term memory (LTM) phase using a different novel object (hM4D(Gi)-CNO: 0.054 ± 0.044; hM4D(Gi)-ss: 0.362 ± 0.049; EGFP-CNO: 0.427 ± 0.049) (Figure 4F, G).

For the exploratory behaviour, we performed the HB test. A pre-test without objects in holes was run before the test phase. In the test phase, identical objects were placed in half of the holes. While head-dipping behaviour in holes without objects did not differ between groups (hM4D(Gi)-CNO: 30.76 ± 2.27; hM4D(Gi)-ss: 35.05 ± 2.45; EGFP-CNO: 35.67 ± 2.16), inhibition of Pthlh cells led to reduced neophilia, as shown by fewer head-dips in holes with objects (hM4D(Gi)-CNO: 26.19 ± 1.66; hM4D(Gi)-ss: 33.00 ± 2.42; EGFP-CNO: 32.13 ± 2.08) (Figure 4H). All groups spent similar time in the center zone (hM4D(Gi)-CNO: 32.63% ± 0.90; hM4D(Gi)-ss: 31.45% ± 0.74; EGFP-CNO: 33.91% ± 0.72), suggesting anxiety levels did not account for the reduced exploratory behaviour (Figure 4J). However, CNO administration appeared to reduce overall activity (hM4D(Gi)-CNO: 1 974 ± 91.30; hM4D(Gi)-ss: 2 301 ± 81.23; EGFP-CNO: 2 249 ± 78.29) (Figure 4I).

Cognitive impairments caused by selective silencing of Pthlh cells in the DS do not appear to be driven by anxiety. In the EPM test, all groups spent similar percentages of time in the open arms (hM4D(Gi)-CNO: 29.59 ± 1.98; hM4D(Gi)-ss: 30.22 ± 1.59; EGFP-CNO: 28.27 ± 3.43) (Supplementary Figure 3A, B) and exhibited similar overall activity (hM4D(Gi)-CNO: 652.1 ± 38.12; hM4D(Gi)-ss: 716.2 ± 48.12; EGFP-CNO: 703.3 ± 43.16) (Supplementary Figure 3C), indicating no significant differences in anxiety-related behaviour.

### 5. Striatal Pthlh interneurons receive predominantly local MSN and long-range thalamic inputs

We performed monosynaptic rabies virus (RABV) tracing using both TVA and the high-specificity variant TVA^66T^ [79], to elucidate inputs to Pthlh interneurons from distant brain regions as well as from local striatal neurons (Figure 5A, B).

**Figure 5.**
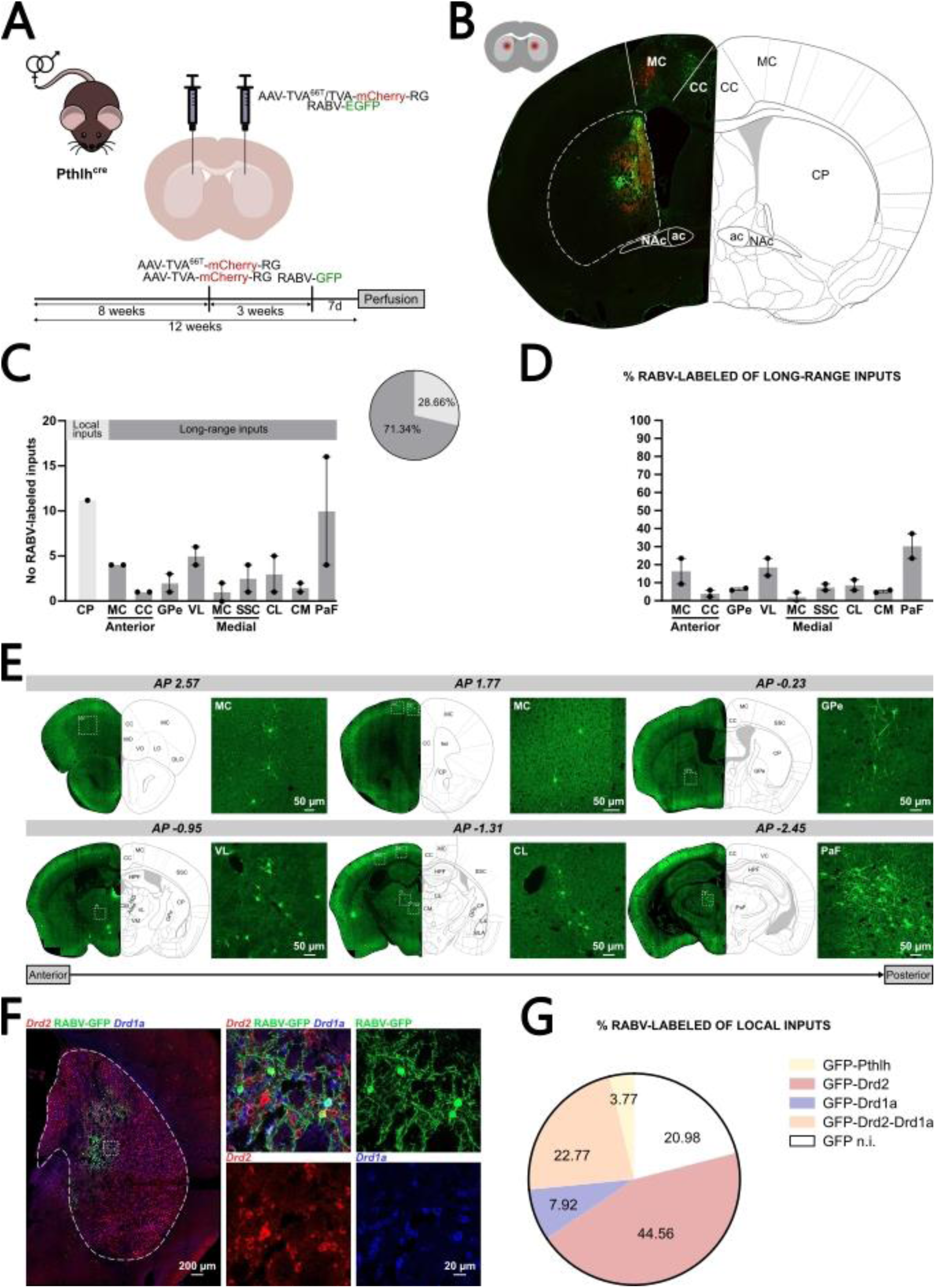
Monosynaptic rabies-based mapping of local and long-range inputs to Pthlh interneurons in the dorsal striatum. **A** Schematic of the viral-genetic strategy used to label monosynaptic inputs to Pthlh interneurons, involving injection of a Cre-dependent helper virus (TVA66T and RG) followed by EnvA-pseudotyped rabies virus (RABV-GFP). **B.** Representative immunofluorescence image showing co-labeled starter cells (tdTomato⁺) and retrogradely labeled input neurons (GFP⁺) within the striatum. **C.** Quantification of GFP⁺ neurons across local (striatal) and long-range input regions, shown as absolute cell numbers and as a percentage of total RABV-labeled inputs. **D.** Proportion of long-range inputs relative to the total monosynaptic inputs to Pthlh interneurons. **E.** Representative images of long-range RABV-labeled inputs from selected regions, including thalamus, cortex, and globus pallidus. **F.** FISH-IHF image showing colocalization of GFP⁺ local inputs with Drd1a and Drd2 mRNA, indicating monosynaptic input from both direct and indirect pathway MSNs. **G.** Proportion of distinct striatal cell types contributing local input to Pthlh interneurons, relative to all RABV-labeled striatal inputs. MC, motor cortex; CC, cingulate cortex; SSC, somatosensory cortex; GPe, external globus pallidus; VL, ventral thalamic nucleus; CM, central medial thalamic nucleus; CL, centrolateral thalamic nucleus; PaF, parafascicular thalamic nucleus.

In our study, 28.66% of the total inputs to striatal Pthlh interneurons originated from local sources (Figure 5C). Given this large local contribution, we next examined the relative input from specific subpopulations. These analyses revealed that most local inputs were medium spiny neurons (MSNs; 75.25%) of different subtypes, including D1 (7.92%), D2 (44.56%), and D1/D2 co-expressing neurons (22.77%). A smaller fraction of inputs arose from other Pthlh interneurons (3.77%) (Figure 5F, G, Supplementary Figure 4A). Long-range inputs to dorsal striatal Pthlh cells included projections from the motor cortex (MC, 18.75%), cingulate cortex (CC, 4.10%), somatosensory cortex (SSC, 7.59%), external globus pallidus (GPe, 6.43%), ventral thalamic nucleus (VL, 18.74%), central medial thalamic nucleus (CM, 5.27%), centrolateral thalamic nucleus (CL, 8.76%), and parafascicular thalamic nucleus (PaF, 30.37%) (Figure 5D, E, Supplementary Figure 4C). Notably, we did not detect rabies-labeled inputs from the basolateral amygdala (BLA), substantia nigra (SN), or ventral tegmental area (VTA) (Figure 5E, Supplementary Figure 4B).

To complement the RABV tracing approach, we additionally employed a non-specific retrograde neurotracer to map afferent projections to the DS. This method confirmed the presence of inputs from thalamic and cortical regions identified through monosynaptic rabies tracing. However, unlike the RABV-based strategy, the retrograde tracer also revealed projections originating from BLA, SN, and VTA (Figure 5E, Supplementary Figure 4B, D). The absence of labeling from these regions in the RABV experiments rules out Pthlh cell involvement in those circuitries that are mainly involved in motor control.

## Discussion

The present study identifies *Pthlh*-expressing interneurons as a major and functionally distinct GABAergic population in the mouse striatum and establishes the Pthlh^cre^ mouse line as a powerful genetic tool for their selective manipulation. By combining high-sensitive FISH, electrophysiological recordings, chemogenetic inhibition, and monosynaptic tracing, we reveal that Pthlh interneurons form a heterogeneous but spatially stable population that critically contributes to cognitive, but not motor, striatal functions. These findings position Pthlh interneurons as key integrators of local and thalamocortical signals and provide new insight into how interneuron diversity supports striatal computations underlying cognition.

We have demonstrated that *Pthlh*-expressing cells constitute one of the largest interneuron populations within the striatum, encompassing the well-known *Pvalb*-expressing interneurons as previously shown in sc/nRNA-seq studies [39, 44, 45, 47]. Importantly, we found that the size of the Pthlh population remains stable across age, sex, and along anteroposterior axis. Although Pthlh interneurons were only recently identified as a molecularly distinct class, *Pvalb*-expressing interneurons are well known and have been extensively studied for their established role in regulating MSNs [80] and in basal ganglia circuitry [81–83]. However, these studies were performed using Pvalb^cre^ mice, which can cover only about 62% of the total Pvalb population and approximately 60% of the Pthlh population [39] underestimating the contribution of *Pthlh*-expressing cells. Our extensive and detailed histological and quantitative analyses demonstrate that the Pthlh^cre^ line effectively targets the Pthlh interneurons and *Pvalb*-expressing cells (Figure 1D and Supplementary Figure 1D), enabling comprehensive access to the full continuum of Pthlh phenotypes and providing the first opportunity to interrogate the Pthlh population *in vivo*. We also observe an increase with age in the *Pvalb* expression within Pthlh population, suggesting that Pthlh cells undergo progressive functional refinement that may tune inhibitory control during the transition from juvenile to adult cognitive behaviors.

Electrophysiological recordings further substantiate the notion of a physiological continuum within the Pthlh population, encompassing both FS and FSL neurons. The existence of these two subclasses indicate that Pthlh interneurons cannot be fully captured by traditional morphological or molecular markers. The ability of Pthlh cells to bridge features of both classical Pvalb FS interneurons and less excitable GABAergic subtypes suggests they serve as flexible modulators of local network synchrony. Until now, studying the entire Pthlh interneuron population required combining multiple mouse lines [39, 48], since relying on a single available line resulted in the exclusion of a substantial subset of *Pthlh*-expressing cells. This new mouse line enables a complete electrophysiological characterization of this interneuron population and provides a valuable tool for future functional studies.

Chemogenetic silencing of Pthlh interneurons in the dorsomedial striatum revealed a specific role in striatum-dependent cognitive performance, without affecting locomotion or anxiety-like behaviour. These results are consistent with emerging evidence that the DS contributes to cognitive processes, including decision-making and memory [84–87]. The absence of motor impairments contrasts with studies reporting that manipulation of Pvalb interneurons affects movement initiation [88]. These functional differences may be attributed to the spatial distribution of *Pvalb* expression within the striatum, which is enriched in ventrolateral regions that receive inputs from motor cortex [39, 89–93]. This anatomical organization supports the interpretation that the cognitive effects observed here arise from disrupted processing of associative or thalamocortical inputs, rather than from alterations in motor circuitry.

Regarding circuitry, our rabies-based mapping revealed that Pthlh interneurons receive dense inputs from thalamic nuclei—including the ventrolateral, mediodorsal, and parafascicular regions—as well as substantial local feedback from MSNs. Importantly, they did not receive inputs from the BLA, SN, or VTA (which were observed with non-specific retrograde tracing of the same striatal injection site), ruling out an involvement of Pthlh interneurons in those circuitries that are primarily associated with motor control. While early works suggested that Pvalb interneurons primarily receive inputs from cortex and globus pallidus, with only minimal thalamic innervation [94–97] other studies have shown that Pvalb interneurons also integrate inputs from intralaminar thalamus, thalamic reticular nucleus and subthalamic nucleus [98, 99]. Our findings position Pthlh interneurons as nodes for integrating multimodal sensory and cognitive information, potentially coordinating striatal output during learning and decision-making. Their strong thalamic innervation resembles that of cholinergic interneurons [35], yet, as inhibitory interneurons, Pthlh cells may instead mediate rapid and temporally precise gating of incoming signals.

Although the present study provides a comprehensive cellular and functional characterization of Pthlh interneurons, several questions remain. Future work combining in vivo calcium imaging or optogenetics with behaviorur will be required to dissect how Pthlh activity patterns encode specific cognitive variables. Furthermore, given the neuroprotective and calcium-regulatory roles of Pthlh signaling [54, 100], exploring whether these interneurons participate in neuroendocrine modulation or contribute to the vulnerability of basal ganglia circuits in disease will be an important direction. Alterations in PTH/PTHLH pathways have been linked to cognitive impairment and mood disorders [64–66], suggesting that Pthlh interneurons might represent a cellular substrate connecting systemic endocrine states with striatal circuit function.

In summary, our findings identify Pthlh interneurons as a major inhibitory class that orchestrates striatal microcircuit activity and cognitive processing. The newly developed Pthlh^cre^ mouse line provides a robust and selective genetic tool to access the entire Pthlh population, overcoming the limitations of previous models that only allowed partial detections. This resource will be instrumental for future mechanistic and translational studies aimed at understanding how inhibitory diversity underlies basal ganglia function and its disruption in neuropsychiatric and neurodegenerative disorders.

## Acknowledgements

The authors acknowledge support from grants PID2022-136526OB-I00by MCIN/AEI /10.13039/ 501100011033 and ERDF/EU (A.B.M.-M.) and RyC programme RYC-2017-22594 (A.B.M.-M.), the Swedish Foundation for Strategic Research FFL 18-0314 (A.B.M.-M.), and the Plan Propio of University of Cádiz (2020-053 / PU / EPIF-FPI-CT / CP to M.D.-S).

We thank the Central Research Services in Health Sciences, and Animal Experimentation Service from the University of Cádiz and Comparative Medicine Biomedicum (KM-B) from Karolinska Institutet.

## AUTHOR CONTRIBUTIONS

M.D.-S. and A.B.M.-M. designed the study; M.D.-S., L.H., M.H., M. Ll.-T., and A.B.M-M carried out experiments; M.D.-S., M.H and A.B.M.-M. performed data analysis; E.B. and J. H.-L provided resources, A.B.M.-M. secured funding; M.D.-S. and A.B.M.-M. wrote the manuscript with comments and reviewing from all authors.

## ETHICS APROVAL

All procedures involving animals were carried out in accordance with the European Directive 2010/63/EU and Spanish Law RD 53/2013 on the protection of animals used for scientific purposes in case of University of Cádiz housing and in accordance with guidelines, permissions and animal protocols approved by Stockholms Djurförsöksetiska Nämnd (6351-2019, 1592-2020, 6211-2021) for Karolinska Institutet colony. Experimental protocols were approved by the Ethics Committee for Animal Experimentation of the School of Medicine, University of Cádiz (Spain) and by the local committees for ethical experiments on laboratory animals (Stockholms Norra Djurförsöksetiska nämnd, Sweden).

