## Supplementary Figures for "A novel key player in cognitive striatal processes: the Pthlh interneurons"

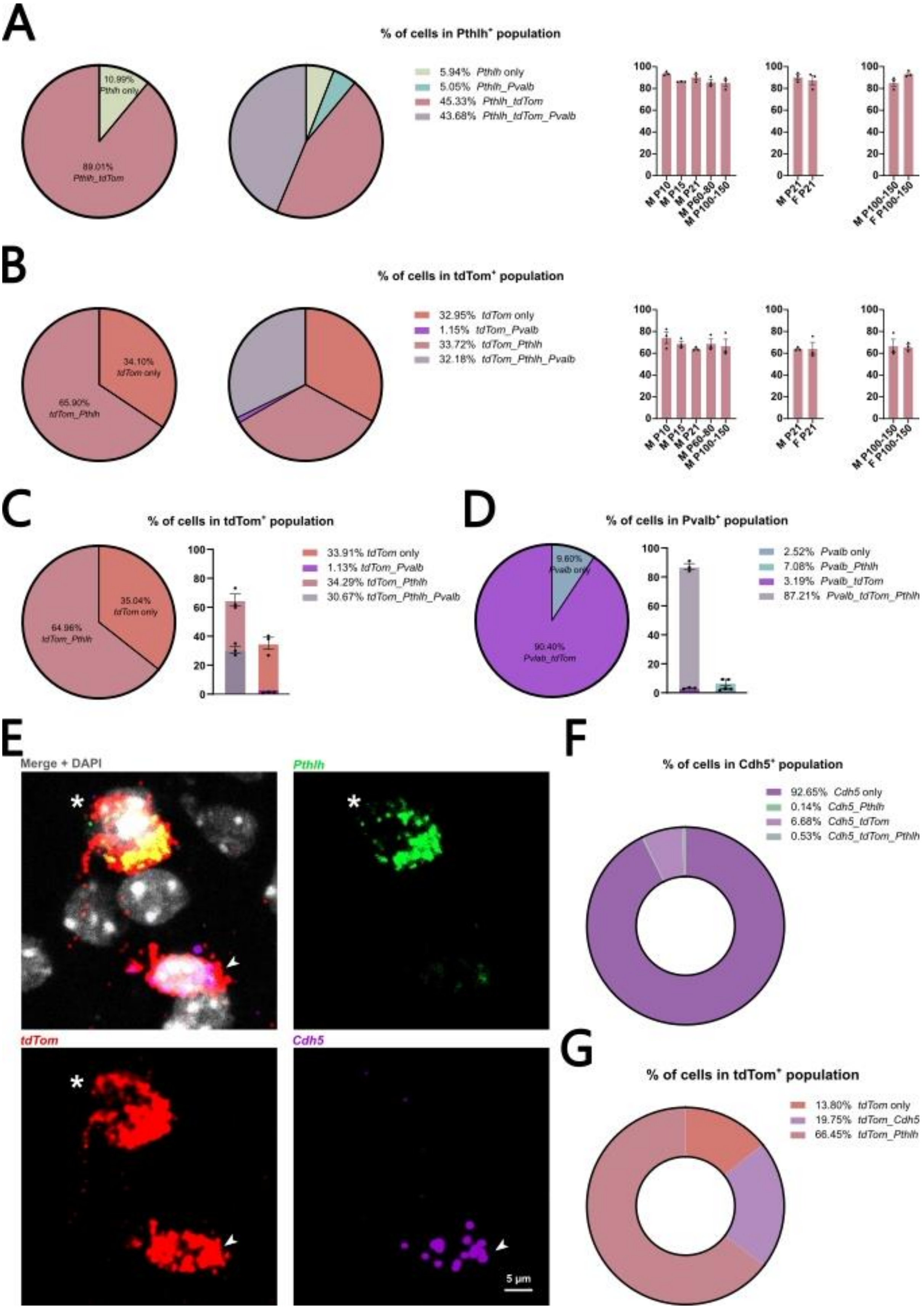


**Supplementary Figure 1. Validation of the Pthlh$^{tdTomato}$ line across different ages and sexes. A.** Pieplots showing the proportions of the *Pthlh* population co-expressing *tdTomato* and *Pvalb* (n = 21), and barplots showing the percentage of *Pthlh* cells labeled by *tdTomato*

with no significant differences across age groups (left), sex at P21 (middle), or sex at P100–150 (right) (N = 3 per group). **B.** Same as in A but analyzing the reporter population (*tdTomato*$^{+}$) co-expressing or not *Pthlh* and *Pvalb*. **C.** Pieplot (left) and barplot (right) showing the distribution of *tdTomato*$^{+}$ cells in P100–150 male mice co-expressing *Pthlh* and *Pvalb* (n = 3). **D.** Same as in C but showing the proportion of *Pvalb*-expressing cells labeled by the reporter. **E.** Fluorescent *in situ* hybridization for *Pthlh*, *tdTomato* and *Cdh5*, asterisk marks a double-positive cell for *Pthlh* and *tdTom* and arrowhead indicates an endothelial cell (*Cdh5*$^{+}$) labeled by the reporter. **F.** Pieplot quantifying the proportion of endothelial cells co-expressing *tdTomato* and *Pthlh* in P100–150 male mice (n = 3). **G.** Same as in F but showing the percentage of the *tdTomato*$^{+}$ population that is either *Pthlh*$^{+}$ or *Cdh5*$^{+}$. Error bars represent mean ± SEM.

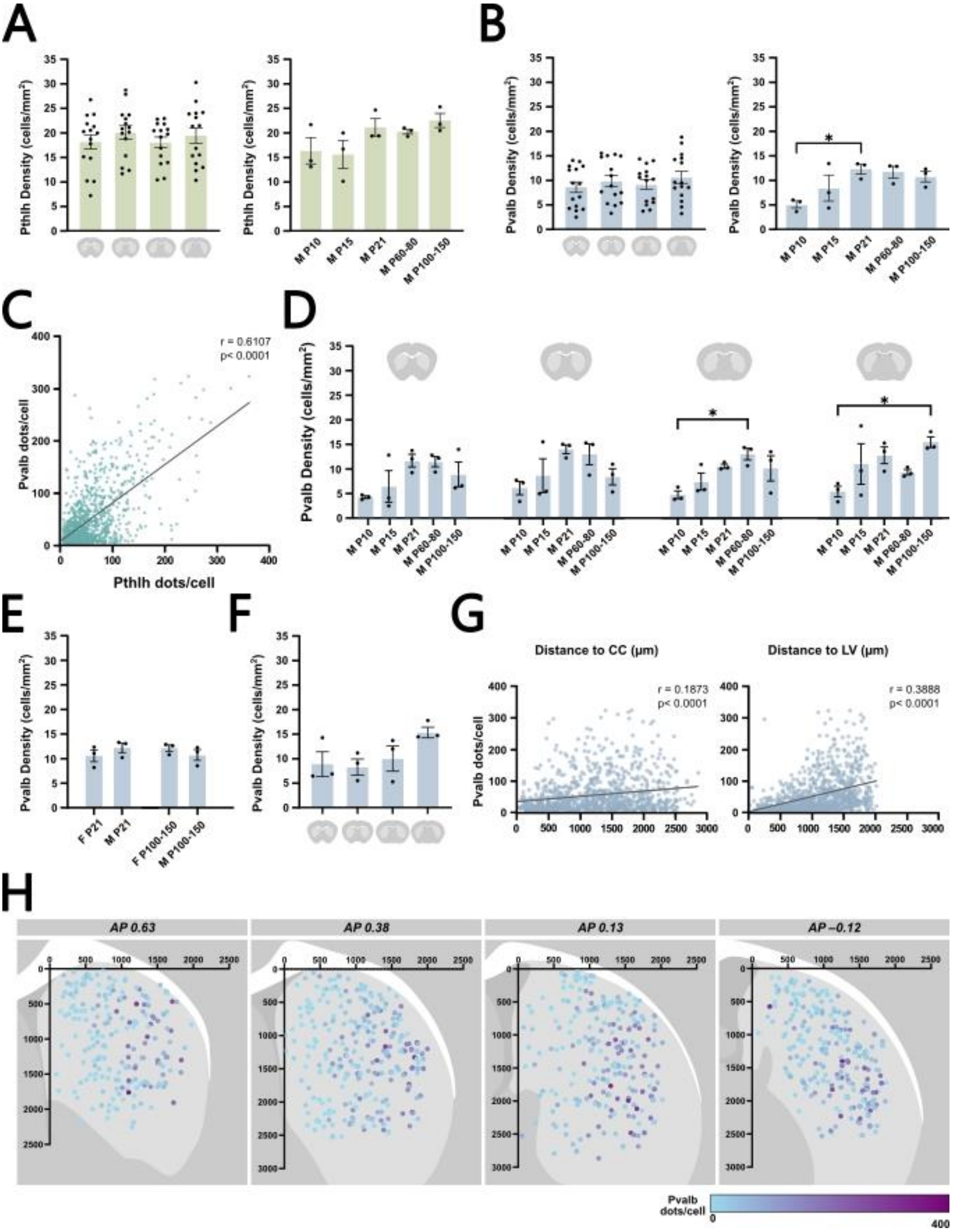


**Supplementary Figure 2. Histological characterization of *Pvalb*-expressing interneurons within the Pthlh population in the dorsal striatum. A.** Barplot showing the density of *Pthlh*-expressing cells along the rostro-caudal axis at four stereotaxic levels: Bregma +0.63 ± 0.05, +0.38 ± 0.05, +0.13 ± 0.05, and –0.12 ± 0.05 mm across all groups (n = 213 per group) (left). Barplot showing the density of *Pthlh*-expressing cells across different experimental male groups (n = 3 per group) (right). **B.** Same as in A, but for *Pvalb*-expressing cells within the Pthlh population. **C.** Scatter plot showing the correlation between *Pthlh* and *Pvalb* expression levels (dots per cell) in P100–150 male mice (n = 3; 1 093 cells). **D.** Barplot showing the density of *Pvalb*-expressing cells across different experimental groups and rostro-caudal levels (n = 3 per group). **E.** Same as in B but comparing females and males at P21 and P100-150 (n = 3 per group). **F.** Barplot showing the rostro-caudal distribution of *Pvalb*-expressing cells in P100–150 male mice (n = 3). **G.** Scatter plots showing the correlation between *Pvalb* expression levels (dots per cell) and anatomical position along the dorsoventral (right) and mediolateral (left) axes in P100–150 male mice (n = 3; 972 cells). **H.** Schematic representation of the spatial distribution and expression levels of *Pvalb*-expressing cells in the dorsal striatum along the rostro-caudal axis at four stereotaxic levels: Bregma +0.63 ± 0.05, +0.38 ± 0.05, +0.13 ± 0.05, and –0.12 ± 0.05 mm (n = 3; 972 cells). Error bars represent mean ± SEM. * $p<0.05$, assessed by one-way ANOVA with repeated measures followed by Tukey's post hoc test. r and p values were acquired using Pearson’s correlation.

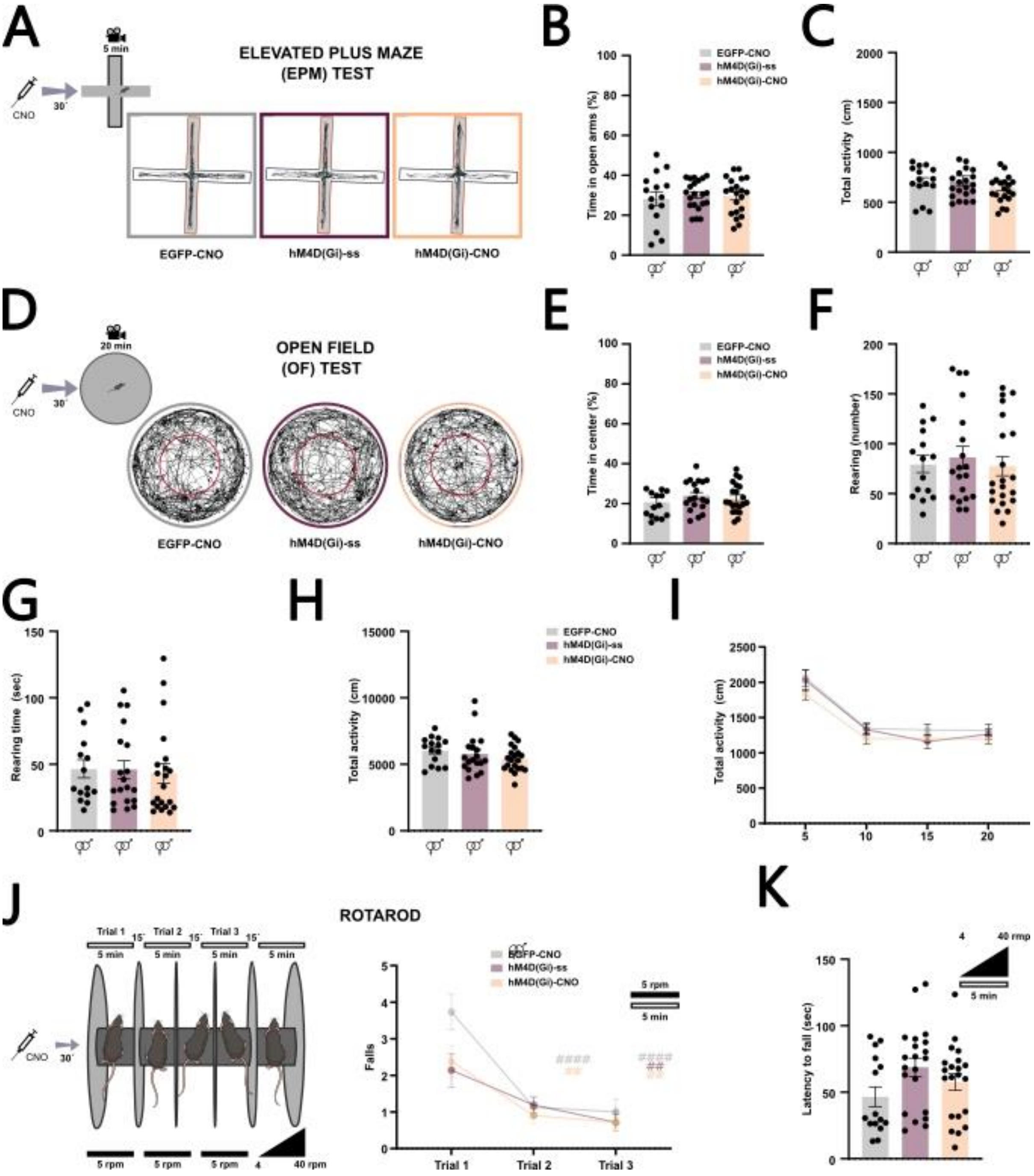


**Supplementary Figure 3. Anxiety-related behavior and motor-skills evaluation following selective inhibition of *Pthlh*-expressing cells. A.** Schematic representation of the Elevated Plus Maze (EPM) test and representative activity traces from each experimental group. **B.** Percentage of time spent in the open arms, shown by group. **C.** Locomotor activity levels by group. **D.** Schematic representation of the Open Field (OF) test and representative activity traces from each experimental group. **E.** Percentage of time spent in the center of the arena, shown by group. **F.** Barplot showing the number of rearings per group. **G.** Time spent doing rearings per group. **H.** Barplot showing locomotor activity levels by group. **I**. Graph representing the activity over consecutive 5-minute intervals shown by group. **J.** Schematic representation of constant speed (5 rpm) and acceleration (4 to 40 rpm) condition performed in rotarod apparatus (left); number of falls in the different trails at constant speed conditions (right). **E.** Latency to fall in the accelerated rotarod by group. Error bars represent mean ± SEM. # p<0.05, ## p<0.01 and #### p<0.0001 versus Trial 1, assessed by one-way ANOVA with repeated measures followed by Tukey's or Kruskal-Wallis post hoc test.

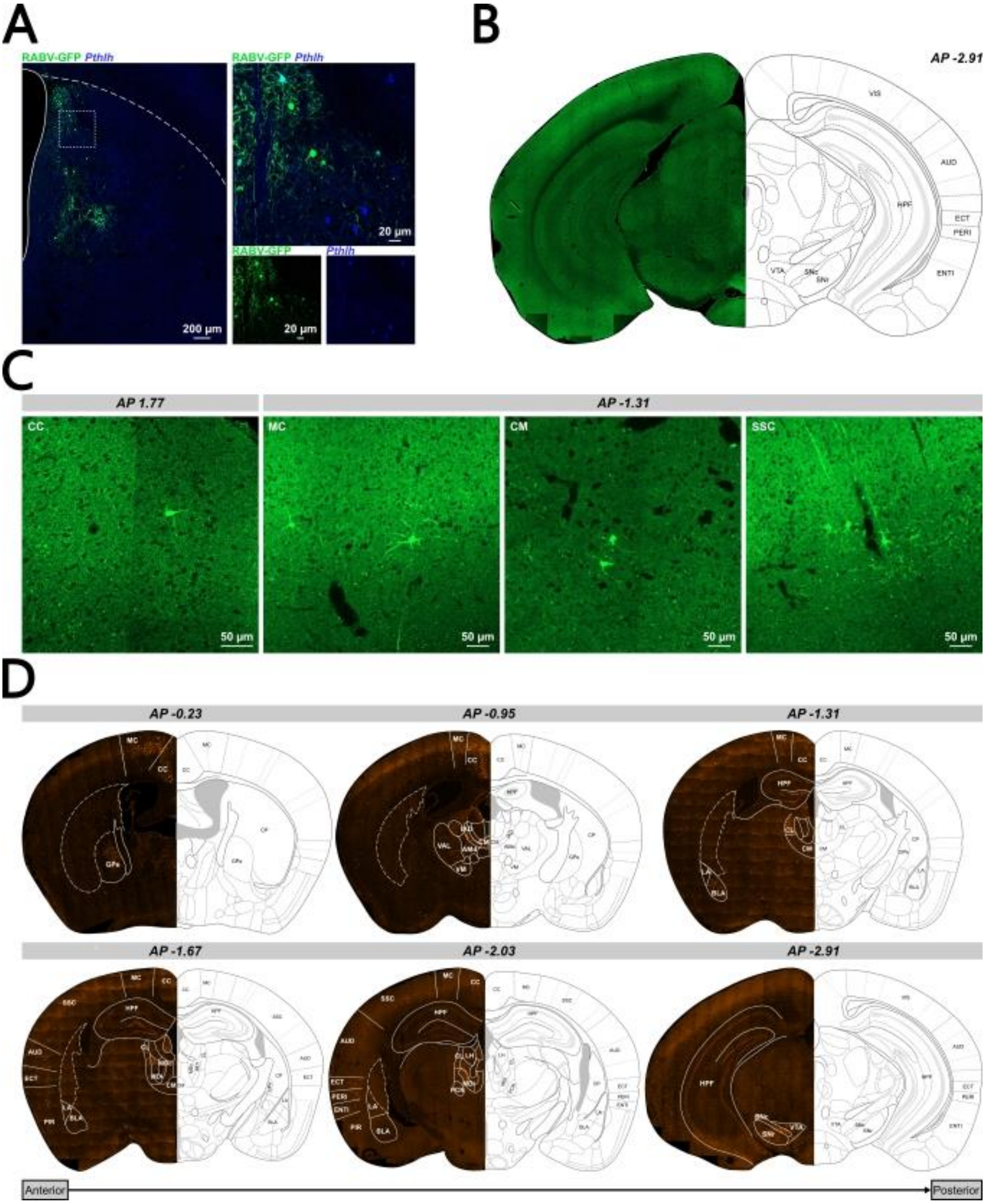


**Supplementary Figure 4. Monosynaptic inputs to Pthlh interneurons in the dorsal striatum. A.** FISH-IHF image showing colocalization of GFP$^{+}$ local inputs with *Pthlh*. **B.** Representative image of long-range RABV-labeled inputs from substantia nigra (SN) and ventral tegmental area (VTA). **C.** Representative images of long-range RABV-labeled inputs from selected regions of AP 1.77 and AP -1.31 brain sections. **D.** Representative images of non-specific neurotracer to striatum from different brain regions. MC, motor cortex; CC, cingulate cortex; SSC, somatosensory cortex; CM, central medial thalamic nucleus; BLA basolateral amygdala.
